# Evolutionary genetic analysis uncovers multiple species with distinct habitat preferences and antibiotic resistance phenotypes in the *Stenotrophomonas maltophilia* complex

**DOI:** 10.1101/138990

**Authors:** Luz Edith Ochoa-Sánchez, Pablo Vinuesa

## Abstract

The genus *Stenotrophomonas* (*Gammaproteobacteria*) has a broad environmental distribution. *S. maltophilia* is its best known species because it is a globally emerging, multidrug-resistant (MDR), opportunistic pathogen. Members of this species are known to display high genetic, ecological and phenotypic diversity, forming the so-called *S. maltophilia* complex (Smc). Heterogeneous resistance and virulence phenotypes have been reported for environmental Smc isolates of diverse ecological origin. We hypothesized that this heterogeneity could be in part due to the potential lumping of several cryptic species in the Smc. Here we used state-of-the-art phylogenetic and population genetics methods to test this hypothesis based on the multilocus dataset available for the genus at pubmlst.org. It was extended with sequences from complete and draft genome sequences to assemble a comprehensive set of reference sequences. This framework was used to analyze 108 environmental isolates obtained in this study from the sediment and water column of four rivers and streams in Central Mexico, affected by contrasting levels of anthropogenic pollution. The aim of the study was to identify species in this collection, defined as genetically cohesive sequence clusters, and to determine the extent of their genetic, ecological and phenotypic differentiation. The multispecies coalescent, coupled with Bayes factor analysis was used to delimit species borders, together with population genetic structure analyses, recombination and gene flow estimates between sequence clusters.

These analyses consistently revealed that the Smc contains at least 5 significantly differentiated lineages: *S. maltophilia* and Smc1 to Smc4. Only *S. maltophilia* was found to be intrinsically MDR, all its members expressing metallo-β-lactamases (MBLs). The other Smc lineages were not MDR and did not express MBLs. We also obtained isolates related to *S. acidaminiphila*, *S. humi* and *S. terrae*. They were significantly more susceptible to antibiotics than *S. maltophilia*. We demonstrate that the sympatric lineages recovered display significantly differentiated habitat preferences, antibiotic resistance profiles and beta-lactamase expression phenotypes, as shown by diverse multivariate analyses and robust univariate statistical tests. We discuss our data in light of current models of bacterial speciation, which fit these data well, stressing the implications of species delimitation in ecological, evolutionary and clinical research.

## 1. Introduction

Bacterial species identification and delimitation are non-trivial tasks, which are critical in certain settings such as the clinic, bio-terrorism and industry. More generally, the conclusions drawn from evolutionary and ecological analyses are strongly dependent on organismal classification, as species are the relevant units of diversity (Vinuesa et al., 2005b;Koeppel et al., 2008;Shapiro et al., 2016). Proper species delimitation is a requisite to discover the species-specific phenotypic attributes underlying their ecological niche differentiation (Cadillo-Quiroz et al., 2012;Shapiro et al., 2012;Cordero and Polz, 2014).

We hypothesized that problems with species delimitations have hindered progress in systematic, taxonomic and ecological research on the ubiquitous genus *Stenotrophomonas* (Gammaproteobacteria, *Xhanthomonadales*, *Xanthomonadaceae*) (Palleroni and Bradbury, 1993), which currently comprises 12 validly described species (http://www.bacterio.net/stenotrophomonas.html). This limitation particularly affects the *S. maltophilia* species complex (Smc) (Svensson-Stadler et al., 2011), which has long been recognized to have a broad ecological distribution, being associated with humans, animals, plants and diverse anthropogenic and natural environments (Berg et al., 1999;Ryan et al., 2009;Berg and Martinez, 2015). Although different genotyping methods, particularly AFLPs (Hauben et al., 1999), rep-PCR (Adamek et al., 2011) and multilocus sequence analysis/typing (MLSA/MLST) (Kaiser et al., 2009;Vasileuskaya-Schulz et al., 2011) have clearly revealed the existence of multiple genomic groups within the Smc, proper recognition of species borders within the complex has not yet been satisfactorily achieved. This has ultimately hindered the discovery of statistically significant associations between species and traits such as habitat preferences, antibiotic resistance phenotypes and pathogenicity potential (Adamek et al., 2011;Berg and Martinez, 2015;Deredjian et al., 2016). *S. maltophilia* is an important globally emerging and multidrug-resistant (MDR) opportunistic pathogen causing difficult-to-treat infections (Chang et al., 2015). High mortality rates are reported mainly in the immunocompromised, cancer and cystic fibrosis patients, as well as those with central venous catheters or long-lasting antibiotic therapy (Looney et al., 2009;Brooke, 2012). Therefore, the identification of significant genotype-phenotype associations is critical for the safe use of particular strains from the Smc with high potential for diverse environmental biotechnologies such as bioremediation, plant growth promotion and protection (Ryan et al., 2009;Berg and Martinez, 2015).

The main objectives of this study were: *i*) to identify genetically cohesive and differentiated sequence clusters (genospecies) among a collection of environmental *Stenotrophomonas* isolates by using a combination of state-of-the-art phylogenetic and population genetic methods; *ii*) to test whether such lineages exhibit distinct phenotypic and ecological attributes, as predicted by current models of bacterial speciation. We used the multilocus dataset for the genus available at pubmlst.org (Kaiser et al., 2001;Vasileuskaya-Schulz et al., 2011), and extended it with sequences extracted from complete (Crossman et al., 2008;Lira et al., 2012;Zhu et al., 2012;Davenport et al., 2014;Vinuesa and Ochoa-Sánchez, 2015) and draft (Patil et al., 2016) genome sequences to assemble a comprehensive MLSA dataset with representative strains of 11 out of 12 validly described *Stenotrophomonas* species. We used this reference dataset to study our collection of environmental *Stenotrophomonas* isolates (*n* = 108) recovered from the sediments and water column of several rivers with contrasting levels of contamination in the state of Morelos, Central Mexico.

For an initial exploration of this dataset, we used thorough maximum-likelihood tree searching. The evidence from this phylogenetic analysis was used to define diverse species border hypotheses, which were formally evaluated in a Bayesian framework under the multispecies coalescent (MSC) model (Rannala and Yang, 2003;Edwards et al., 2007;Degnan and Rosenberg, 2009) by subjecting them to Bayes factor (BF) analysis (Kass and Raftery, 1995). To the best of our knowledge, this is the first study that evaluates the utility of this Bayesian statistical framework for bacterial species delimitation, which is emerging as a successful and promising strategy for species delimitation in plants and animals (Fujita et al., 2012;Aydin et al., 2014;Grummer et al., 2014). The MSC model is independent of gene concatenation and acknowledges the very well known fact that gene trees have independent evolutionary histories embedded within a shared species tree (Degnan and Rosenberg, 2006;Rosenberg, 2013). The basic MSC model assumes that gene tree discordance is solely the result of stochastic coalescence of gene lineages within a species phylogeny. Populations, rather than alleles sampled from single individuals, are the units to infer phylogeny in the MSC framework, effectively connecting traditional phylogenetic inference with population genetics, providing estimates of topology, divergence times and population sizes (Rannala and Yang, 2003;Edwards et al., 2007;Heled and Drummond, 2010).

Current microbial speciation models predict that bacterial species-like lineages should be identifiable by significantly reduced gene flow between them, even when recombination levels are high within species. Such lineages should also display differentiated ecological niches (Koeppel et al., 2008;Cadillo-Quiroz et al., 2012;Shapiro and Polz, 2014). This study shows the power of modern phylogenetic and population genetic methods to delimit species borders in bacteria and demonstrates that the Smc, as currently defined in pubmlst.org, genome databases and literature, contains multiple genospecies that are ecologically and phenotypically differentiated. We discuss our findings and approaches in the light of current models of bacterial speciation, highlighting the practical implications and ecological relevance of proper species delimitation.

## 2. Materials and Methods

### Sampling sites and isolation of environmental *Stenotrophomonas* strains

*Stenotrophomonas* strains were recovered from the sediments and water columns at 6 sites of 4 rivers and streams in the State of Morelos, Central Mexico (Table 1, supplementary Fig. S1). These sites experience different levels of anthropogenic pollution, broadly classified as low (L), intermediate (I) and high (H), based on triplicate counts of thermo-tolerant coliforms (TTCs) on mFC agar (Oxoid). Thermotolerant *E. coli* (TTEc) counts were obtained on modified m-TEC agar (USEPA, 2002), using the one-step membrane filtration (0.45 µm) method (APHA, 2005). Water samples were taken in sterilized 1L recipients at 5–20 cm depth (2 per site). Six physico-chemical parameters of the water columns were measured using a HANNA multi-parametric HI9828 instrument operated in continuous measurement mode, for 1 minute, along a 10 m transect (Fig. S2). Sediment samples (3 per site, along a 3 m linear transect) were taken from the same sites in sterile plastic cores dug 2–3 cm deep into the sediment. Samples were kept on ice until processing within 4–8 hrs (APHA, 2005). Sampling took place at the end of the dry season (April-May), between 2012 and 2014. Oligotrophic [NAA (Aagot et al., 2001) and R2A (Ultee et al., 2004) agar] and rich media [LAM (Jawad et al., 1994) and MacConkey], supplemented or not with antibiotics [trimethoprim 30 + carbenicillin 100, ciprofloxacin 4, ceftazidime 8, cefotaxime 4, imipenem 4 (µg/ml)] were used to isolate bacteria from these samples by plating 100 µl of serial dilutions (1 to 10e^−4^) in triplicate for each sample and incubation at 30°C for up to 24 hrs. Single colonies were repeatedly streaked on the same media for strain purification. Bacteria were routinely grown on LB and stored frozen in this medium supplemented with 20% (V/V) glycerol at −80°C.

**Table 1.**
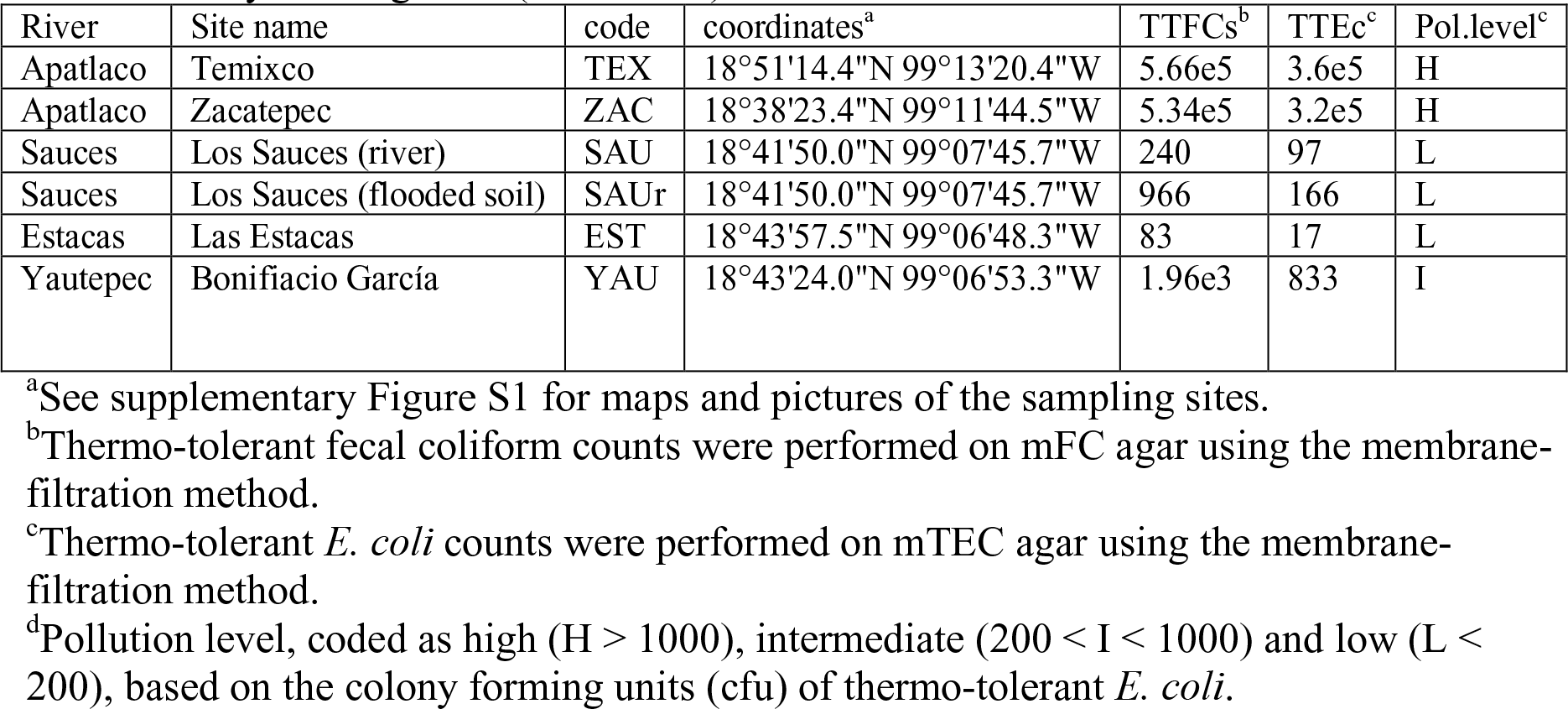
Sampling sites for this study in Morelos, Mexico and pollution level based on counts of thermotolerant fecal coliforms (TTFCs) and thermotolerant *E. coli* (TTEc) colony forming units (cfu/100 ml) measured in the water column.

### Determination of antibiotic resistance and β-lactamase expression profiles

A total of 15 antimicrobials from 6 families and two inhibitor/β-lactamase combinations were used to determine the resistance profiles of each strain by streaking them in parallel on agar plates supplemented with the antibiotics and concentrations indicated in supplementary Table S1. Double disk synergism (DDS) assays were performed to determine the expression phenotypes of specific β-lactamase types [Ambler class A extended spectrum beta-lactamases (ESBLs), class B metallo-beta-lactamases (MBLs) and class C cephalosporinases (AmpC)], as detailed in the legend to Fig S10. The antibiotic breakpoint concentrations and growth inhibition zones were interpreted according to the 26^th^ edition of the Clinical and Laboratory Standards Institute (CLSI, 2016) values for *Stenotrophomonas*, *Pseudomonas aeruginosa* or *Enterobacteriaceae*, when not available for the first or second genus, respectively (cutoff values are shown in Tables S1 and S2).

### PCR amplification of 16S rDNA sequences and their phylogenetic analysis

All strains recovered were classified at the genus level by phylogenetic analysis of the 16S rRNA gene (*rrs*) sequences amplified with the universal fD1/rD1 primers (Weisburg et al., 1991), as previously described (Vinuesa et al., 2005a), and detailed in the supplementary material (supplementary protocol 1).

### PCR amplification and maximum likelihood phylogenetic analysis of multilocus sequence data

For multilocus sequence analysis (MLSA) of environmental *Stenotrophomonas* isolates we used the primers and conditions reported at http://pubmlst.org/smaltophilia/, except for the mutM_steno_6F (5’-ytdcccgaagtmgaaacyac-3’) and mutM_steno_684R (5’-gcagytcctgytcgaartarcc-3’) primers, which were designed *de novo* using the Primers4Clades server (Contreras-Moreira et al., 2009) fed with *mutM* orthologues identified using the GET_HOMOLOGUES package (Contreras-Moreira and Vinuesa, 2013) from *Stenotrophomonas* genome sequences (data not shown). PCR amplicons were purified and commercially sequenced at both strands by Macrogen, (South Korea). Raw reads were assembled with the phredphrap script (de la Bastide and McCombie, 2007), codon-based multiple sequence alignments generated with an in-house Perl script, and MSA borders stripped to match the reference pubmlst.org profiles. Individual gene alignments were concatenated and the resulting matrix subjected to model selection with jModelTest2 (Darriba et al., 2012) for phylogenetic analysis under the maximum likelihood criterion in PhyML3 (Guindon et al., 2010). Tree searches were initiated from 1000 random seed trees and a BioNj phylogeny, under the BEST moves option, as previously described (Vinuesa et al., 2008).

### Identification of the 7 MLSA loci in genome sequences retrieved from GenBank

We selected 24 complete (Crossman et al., 2008;Lira et al., 2012;Zhu et al., 2012;Davenport et al., 2014;Vinuesa and Ochoa-Sánchez, 2015) and draft (Patil et al., 2016) genome sequences available in GenBank to expand our dataset with additional key reference strains. The orthologs of the seven MLSA loci were identified using from single-copy homologous gene clusters computed with the GET_HOMOLOGUES package (Contreras-Moreira and Vinuesa, 2013). We found that the *gap* gene in the draft genome sequence of strain *S. ginsengisoli* DSMC24757^T^ (Acc. No. LDJM00000000) (Patil et al., 2016) contains a thymidine insertion at position 602 that causes a frame-shift mutation and a premature end of the gene. Consequently, the last 72 sites of this sequence were re-coded as ‘?’ (missing characters).

### Sequence data availability

The sequences generated in this study for multilocus sequence analysis were deposited in GenBank under accession numbers KX895367-KX896038.

### Bayesian species delimitation using the multispecies coalescent (MSC) and Bayes factors (BFs)

Bayesian species delimitation from multilocus data under the MSC model was performed using the recent *BEAST2 module (version 0.7.2) for BEAST 2 (Heled and Drummond, 2010;Bouckaert et al., 2014), to evaluate a set of explicit hypotheses of species-boundaries. *BEAST2 was run using the best fitting partitioning scheme (see supplementary protocol 2) and the TrN+G model with empirical frequencies, without rate estimation. Trees were unlinked across partitions, setting the ploidy level to 1 for each gene tree and assuming a constant population IO population model. A non-correlated relaxed log-normal clock (Drummond et al., 2006) was assumed for each partition, fixing the clock rate of the first partition and estimating the remaining ones. A non-calibrated Yule prior was set on the species tree. The default 1/x population mean size prior was changed for a proper inverse gamma prior (Baele et al., 2013), with shape parameter alpha = 2 and scale parameter beta = 2 and an initial value of 0.05. The upper and lower bounds were set to 0.001 and 1000.0, respectively. Path sampling was used to estimate the marginal likelihoods of each species delimitation model in *BEAST2 runs for BF calculations (Lartillot and Philippe, 2006;Baele et al., 2012;Grummer et al., 2014), with the MODEL_SELECTION 1.3.1 package. Each *BEAST2 chain was run for 10^8^ generations, sampling the posterior every 20000^th^, with 10 replicate runs and the alpha value set to 0.3, applying 50% burnin. A final triplicate *BEAST2 analysis was set up to get the final estimate of the multispecies phylogeny under the best delimitation model with the same parameters, priors, chain length and sampling frequency described above. Convergence and mixing of replicate runs was checked in tracer (http://tree.bio.ed.ac.uk/software/tracer/), as well as the effective sample size values for each parameter. The species tree corresponding to the best species-delimitation hypothesis was visualized with densitree (Bouckaert, 2010), on combined post-burnin (50%) species tree files generated with logcombiner. A summary tree was generated from the latter with treeannotator and visualized with FigTree v1.4.2 http://tree.bio.ed.ac.uk/software/figtree/.

### DNA polymorphism, population structure and recombination analyses

Descriptive statistics for DNA polymorphisms, population differentiation, gene flow, diverse neutrality and population growth tests, as well as coalescent simulations, were computed with DNAsp v.5.10.01 (Rozas et al., 2003), as previously described (Vinuesa et al., 2005b). Bayesian analysis of population structure based on multilocus sequence data was performed in STRUCTURE v2.3.4 under the admixture and correlated gene frequencies models (Pritchard et al., 2000;Falush et al., 2003;2007). Twenty runs were launched for each *K* value between 2 and 10, with 10^5^ steps sampled after a burnin of 2x10^5^ chain iterations. The best *K* value was defined by the Evanno (Evanno et al., 2005) and Pritchard (Pritchard et al., 2000) methods, as implemented in CLUMPAK (Kopelman et al., 2015). Estimation of recombination rates of selected lineages was performed with ClonalFrameML v1.0.9 (Didelot and Wilson, 2015), using ML trees and Ti/Tv ratios estimated under the HKY85+G model with PhyML3 (Guindon et al., 2010).

### Statistical analyses

All statistical and graphical analyses were performed with base R (R Development Core Team, 2016) and add-on packages. Basic data manipulation, transformation and graphical displays were achieved with functions of the tidyverse metapackage (https://CRAN.R-project.org/package=tidyverse). Tests for normality, homoscedasticity, outliers and skew were performed with the car (https://cran.r-project.org/package=car) and moments (https://cran.r-project.org/package=moments) packages. Robust ANOVA and associated *post-hoc* analyses (Wilcox, 2016) were performed with the WRS2 package (https://cran.r-project.org/package=WRS2). Empirical distributions of test statistics were generated by bootstrapping with the boot package (https://cran.r-project.org/package=boot). Multivariate association plots for categorical data were performed with the vcd package (https://cran.r-project.org/package=vcd). Multiple correspondence analysis (MCA) was performed with the FactoMineR (https://cran.r-project.org/package=FactoMineR) and factoextra packages (https://cran.r-project.org/package=factoextra).

## 3. Results

### 3.1 Evaluation of different isolation media for the recovery of *Stenotrophomonas* from aquatic ecosystems with contrasting degrees of fecal contamination

We sampled 6 sites located in four rivers/streams of Morelos (supplementary figure S1) that were ranked into three categories based on their pollution level (low, intermediate, high), based on counts of thermotolerant fecal coliforms and *E. coli* (Table 1). Additional physicochemical parameters of each sampling site are presented in Fig. S2. Classification of the isolates at the genus level was based on the phylogenetic analysis of 16S rRNA gene sequences (*n* = 697), as shown in Fig. S3. *Stenotrophomonas* was the second most abundant genus (*n* = 154, 22.1 %) recovered in our collection, after *Pseudomonas* (*n* = 239, 34.3 %), as shown in Fig S4. The inset in Fig. S4 shows a Trellis barplot summarizing the relative efficiency of the different microbiological media tested for the recovery of *Stenotrophomonas*. The analysis reveals that environmental *Stenotrophomonas* strains can be efficiently recovered on the oligotrophic NAA medium supplemented with imipenem (8 µg/ml; > 90% recovery efficiency). The rich McConkey medium amended with imipenem is also useful (4 µg/ml; ~60%), selecting non-fermenting (whitish) colonies.

### 3.2 Phylogenetic structure of the genus *Stenotrophomonas* and the definition of species border hypotheses

We used intense maximum likelihood (ML) tree searching (see methods) to obtain a global hypothesis of the phylogenetic structure of the genus based on 194 non-redundant multilocus STs (Fig. 1). This dataset contains sequences retrieved from pubmlst.org (seven loci for 103 STs representative of all the “classic” genogroups and clusters defined in previous works (Kaiser et al., 2009;Vasileuskaya-Schulz et al., 2011), plus 24 selected reference strains from different species for which the genome sequences were retrieved from GenBank and the 7 loci extracted, as explained in methods. This comprehensive set of reference strains comprises 11 out of 12 validly describes species of the genus as of April 2017. Recently *S. tumulicola* (Handa et al., 2016) was added to the list (www.bacterio.net/stenotrophomonas.html; last access April 10th, 2017), which is the only species missing from our analysis, as it lacks a genome sequence or MLSA data in pubmlst.org. To this set we added the sequences generated in this study for 108 environmental isolates from Mexican rivers, comprising 63 haplotypes (distinct multilocus sequence types). Figure 1A presents the best hypothesis found among 1001 independent tree searches (the ln*L* profile of the tree search is shown in figure S5A), displaying the Smc clade in collapsed form. The ML tree was rooted using four *Xanthomonas* species as the outgroup, chosen based on the evidence of a comprehensive ML phylogeny of nearly full-length 16S rRNA gene sequences for all type strains currently described in the order *Xanthomonadales* (Fig. S6). *S. panacihumi* (Yi et al., 2010) represents the most basal lineage of the genus, which is consistent with its position on the 16S rRNA gene phylogeny (Fig. S6). However, this species was described based on the *rrs* sequence analysis of a single isolate and currently has a non-validated taxonomic status. Two large clades follow, labeled I and II on figure 1A. All species grouped in clade I are of diverse environmental origin, lacking strains reported as opportunistic pathogens. Three of our environmental isolates tightly cluster with the type strain of *S. acidaminiphila* (Assih et al., 2002), including strain ZAC14D2_NAIMI4_2, for which we have recently reported its complete genome sequence (Vinuesa and Ochoa-Sánchez, 2015). The type strain of this species (AMX19^T^, LMG22073^T^) was isolated from the sludge of a lab-scale anaerobic chemical waste water reactor in Iztapalapa, Mexico City, 1999 (Assih et al., 2002). A single isolate of our collection is phylogenetically related to *S. humi*, while 7 others form a perfectly supported clade with the type strain of *S. terrae* (Heylen et al., 2007). In conclusion, all environmental isolates grouped in clade I were conservatively classified as indicated by the labeled boxes on Fig. 1A, based on the very strong support of the monophyletic clusters formed with the corresponding type strains.

**Figure 1.**
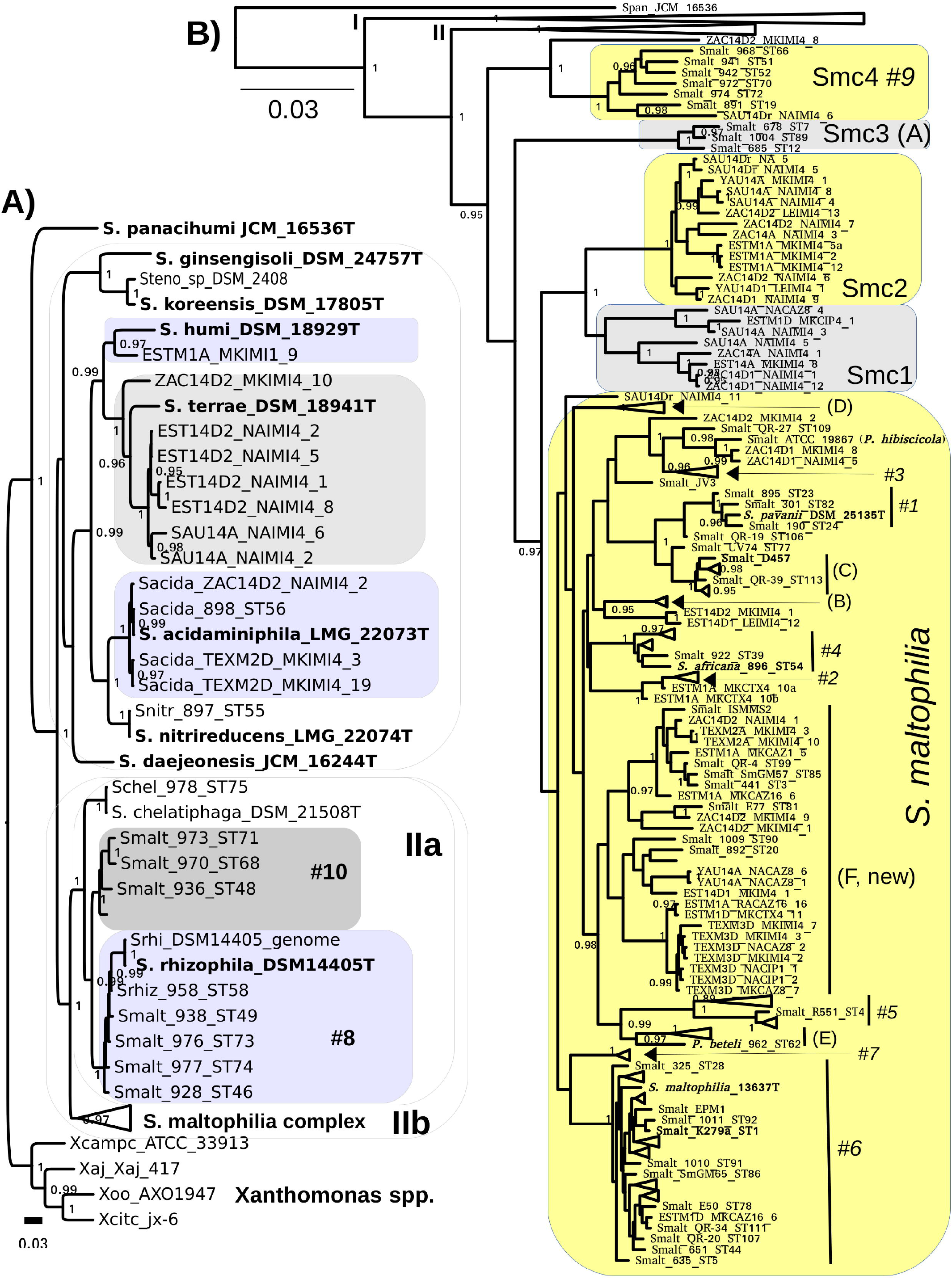
Maximum likelihood multilocus phylogeny of the genus *Stenotrophomonas* and species delimitation hypotheses. The tree shown corresponds to the best one found out of 1001 searches under the GTR+G model and BEST moves, using the concatenated alignment (7-loci) for 194 non-redundant STs, containing all validly described species of the genus except for *S. tumulicola*, additional key reference strains and 63 haplotypes from the 108 Mexican environmental isolates analyzed in this study. A) The S. maltophilia complex (Smc) clade is collapsed in this tree. B) The Smc clade is displayed, collapsing the clades visible in B. Clades are labeled using the A-E codes of Kaiser et al. (2009) and #1-#10 of Hauben et al. (1999) for easy cross-reference. The shaded areas indicate species assignation hypotheses specifically evaluated in this study by the multispecies coalescent using Bayes factors and by population genetics analyses (Tables 2 and 3). The combined evidence of both approaches reveals that the Smc complex should be split into S. maltophilia and four new species lineages: Smc1, Smc2, Smc3 and Smc4 (Fig. 1B). We also show (Fig 1A) that genotype #10 represents a non-described species that should not be merged with *S. rhizophila* clade #8 (Table 2). Type and other key reference strains are highlighted in bold-face. The bar indicates the number of expected substitutions per site under the GTR+G model.

Clade IIa groups strains of *S. chelatiphaga* and *S. rhizophila* and clade IIb groups strains of the *S. maltophilia* species complex (Smc) (Fig. 1A). Both of them hold human opportunistic pathogens and environmental isolates, and suffer from taxonomic problems. The taxonomic inconsistency of classifying strains of genogroups #8 and # 10 as *S. maltophilia* was previously recognized (Vasileuskaya-Schulz et al., 2011). They cluster within the *S. chelatiphaga-S. rhizophila* clade, making *S. maltophilia* polyphyletic. *S. maltophilia* strains of genogroup #8 were already recognized to belong to *S. rhizophila*, but the taxonomic status of its sister clade, genogroup #10, holding strains labeled as *S. maltophilia*, was not clarified (Vasileuskaya-Schulz et al., 2011).

Figure 1B shows the same phylogeny presented in Fig. 1A, but collapsing clades I and IIa, and displaying the Smc strains grouped in cluster IIb. All terminal clades containing only reference sequences were also collapsed to avoid excessive cluttering. Figure S5B displays the same tree but without collapsing those clusters. The Smc clade was conservatively split into 5 potential species (labeled boxes in Fig. 1B) based on the deep and strongly supported branches subtending MLSA phylogroups Smc1-Smc4, and taking into account the location of the well characterized *S. maltophilia* model strains K279a (Crossman et al., 2008), D457 (Lira et al., 2012) and the type strain of the species ATCC 13637^T^(Davenport et al., 2014). The latter three are spread across the clade labeled as *S. maltophilia* (Fig. 1B). Each of the phylogroups Smc1 to Smc4 currently holds ecologically coherent groups of strains: Smc1 and Smc2 contain exclusively Mexican river isolates recovered in this study, Smc3 groups cystic fibrosis isolates and Smc4 predominately rhizosphere isolates from diverse plants and parts of the world. In contrast, the large *S. maltophilia* clade holds heterogeneous groups of isolates of clinical and environmental origin, including the previously defined MLSA phylogroups A to E (Kaiser et al., 2001;Vasileuskaya-Schulz et al., 2011) and AFLP genogroups #1 to #10 (Hauben et al., 1999). We conservatively define a new MLSA phylogroup F, comprising isolates of clinical and environmental origin from different continents, but dominated by river isolates reported herein (Fig. 1B). We note that *S. pavanii* DSM 25135^T^, an endophytic N_2_-fixing bacterium isolated from sugarcane in Brazil (Ramos et al., 2010), is clearly nested within genogroup #1, closely related to the reference strain *S. maltophilia* D457 (Fig. 1B). This represents an additional taxonomic inconsistency for the *S. maltophilia* clade not previously reported. Here we suggest that *S. pavanii* is a late heteronym of *S. maltophilia*. *S. africana*, nested within genogroup #4, had already been described as a later heterotypic synonym of *S. maltophilia* (Coenye et al., 2004;Kaiser et al., 2009), the same as *Pseudomonas beteli* 962^T^, related to MLSA genogroup E, and *P. hibiscicola* ATCC19867, related to genogroup #3 (Hauben et al., 1999;Vasileuskaya-Schulz et al., 2011). The latter two species were already recognized in 1990 to be misclassified based on the analysis of an extensive set of phenotypic features, being synonyms of *S. maltophilia* (Van Den Mooter and Swings, 1990). The taxonomic status of *S. tumulicola* (Handa et al., 2016) could not be revised in the present work because it lacks multilocus sequence data.

### 3.3 Bayesian species delimitation based on the multispecies coalescent (MSC) model and Bayes factor (BF) analysis of marginal likelihoods

We used a recent software implementation of the MSC model (Heled and Drummond, 2010;Bouckaert et al., 2014) to test the explicit species delimitation hypotheses highlighted in Figs. 1A and 1B by means of BF analysis of marginal likelihoods (Grummer et al., 2014) in a formal Bayesian statistical framework. The best partitioning scheme (see supplementary protocol 2 and Fig. S7) was used for all *BEAST2 runs. Of particular interest to this work was the evaluation of the following five species delimitation hypotheses within the Smc: split1 (species assignations as defined by the shaded areas on Figs. 1A and 1B), lump_*S maltophilia*+Smc1, lump_*S. maltophilia*+Smc12, lump_*S. maltophilia*+Smc123, lump_*S. maltophilia*+Smc1234, which successively lump the *S. maltophilia* sequence cluster with the proposed Smc1-Smc4 genospecies (Table 2 and Fig. 1B). Analysis of the logfiles for the path sampling runs for each model and replicate had effective sample size (ESS) values > 150 for all parameters, most of them with ESSs >> 500. As shown in Table 2, the split1 hypothesis was the favored one as it attained the highest marginal likelihood. The BF analysis provides overwhelming evidence (Table 2) that the Smc, as actually defined in pubmlst.org, lumps multiple cryptic species, strongly supporting the species delimitation hypothesis presented in Fig. 2B, which conservatively splits the complex into the 5 species: *S. maltophilia* and the four new lineages Smc1, Smc2, Smc3 and Smc4. Smc1 and Smc2 contain only Mexican representatives of environmental *Stenotrophomonas* sampled in this study. The split1 vs. lump_cluster#8+cluster#10 (Fig 1A) model was also evaluated, providing overwhelming evidence in favor of separating the two genogroups #8 and #10 as distinct species (ln-BF > 5, Table 2). Supplementary figures S8 and S9 show the consensus and DensiTree (see methods) species tree representations, respectively, of the merged (3 replicate runs), post-burnin (50%) samples, for the best hypothesis.

**Table 2.**
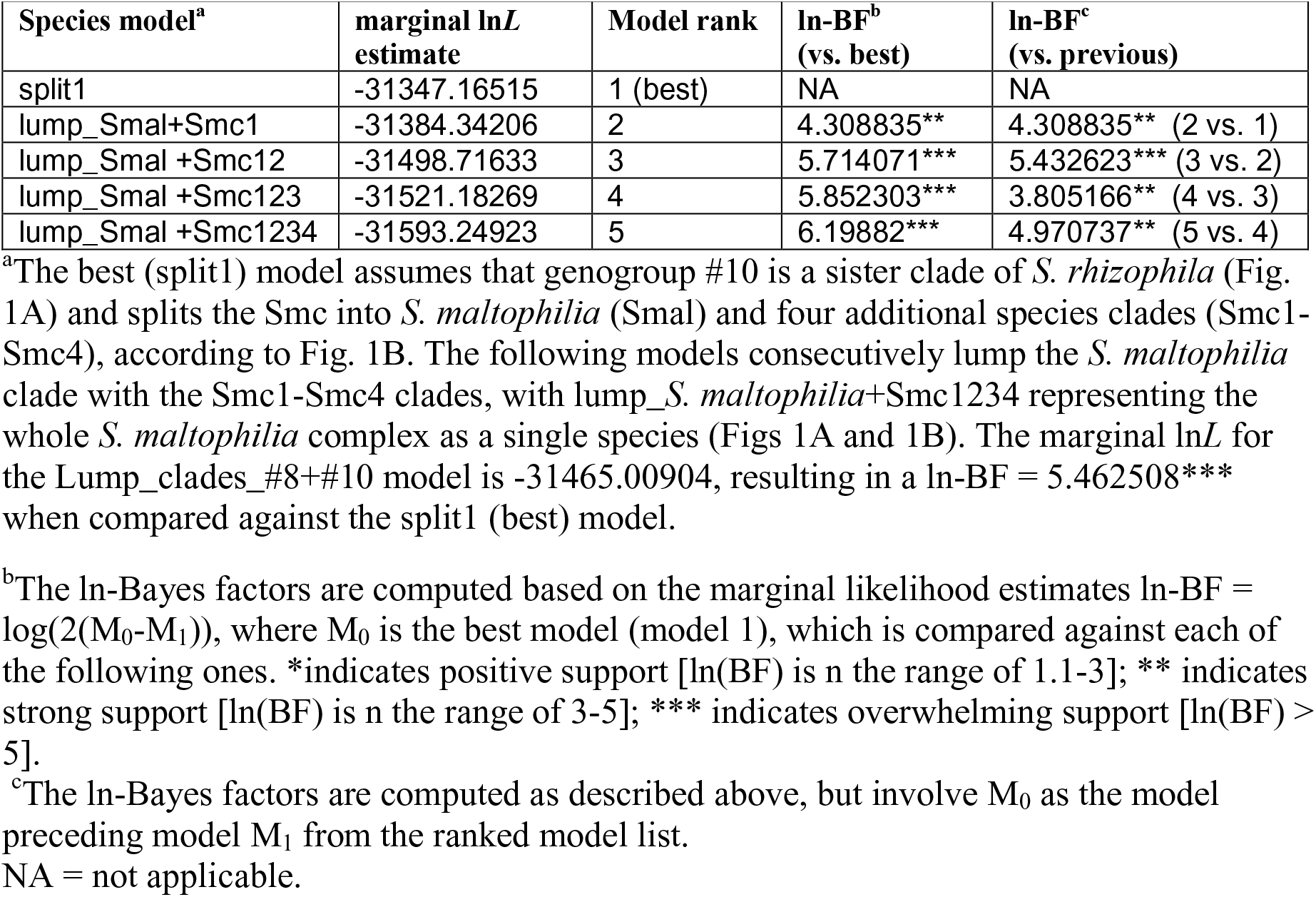
Bayes factor (BF) analysis for 5 species delimitation hypotheses within the *S. maltophilia* complex, plus genogroups #8 and #10, based on marginal likelihoods computed for each hypothesis by path sampling (see methods).

| Species model<sup>a</sup> | marginal lnL estimate | Model rank | ln-BF<sup>b</sup> (vs. best) | ln-BF<sup>c</sup> (vs. previous) |
| --- | --- | --- | --- | --- |
| split1 | -31347.16515 | 1 (best) | NA | NA |
| lump_Smal+Smc1 | -31384.34206 | 2 | 4.308835** | 4.308835** (2 vs. 1) |
| lump_Smal+Smc12 | -31498.71633 | 3 | 5.714071*** | 5.432623*** (3 vs. 2) |
| lump_Smal+Smc123 | -31521.18269 | 4 | 5.852303*** | 3.805166** (4 vs. 3) |
| lump_Smal+Smc1234 | -31593.24923 | 5 | 6.19882*** | 4.970737** (5 vs. 4) |

<sup>a</sup>The best (split1) model assumes that genogroup #10 is a sister clade of *S. rhizophila* (Fig. 1A) and splits the Smc into *S. maltophilia* (Smal) and four additional species clades (Smc1-Smc4), according to Fig. 1B. The following models consecutively lump the *S. maltophilia* clade with the Smc1-Smc4 clades, with lump\_ *S. maltophilia*+Smc1234 representing the whole *S. maltophilia* complex as a single species (Figs 1A and 1B). The marginal lnL for the Lump\_clades\_#8+#10 model is -31465.00904, resulting in a ln-BF = 5.462508\*\*\* when compared against the split1 (best) model.
<sup>b</sup>The ln-Bayes factors are computed based on the marginal likelihood estimates $\ln\text{-BF} = \log(2(M_0 - M_1))$ , where $M_0$ is the best model (model 1), which is compared against each of the following ones. \*indicates positive support [ $\ln(\text{BF})$ is in the range of 1.1-3]; \*\* indicates strong support [ $\ln(\text{BF})$ is in the range of 3-5]; \*\*\* indicates overwhelming support [ $\ln(\text{BF}) > 5$ ].
<sup>c</sup>The ln-Bayes factors are computed as described above, but involve $M_0$ as the model preceding model $M_1$ from the ranked model list.
NA = not applicable.

**Figure 2.**
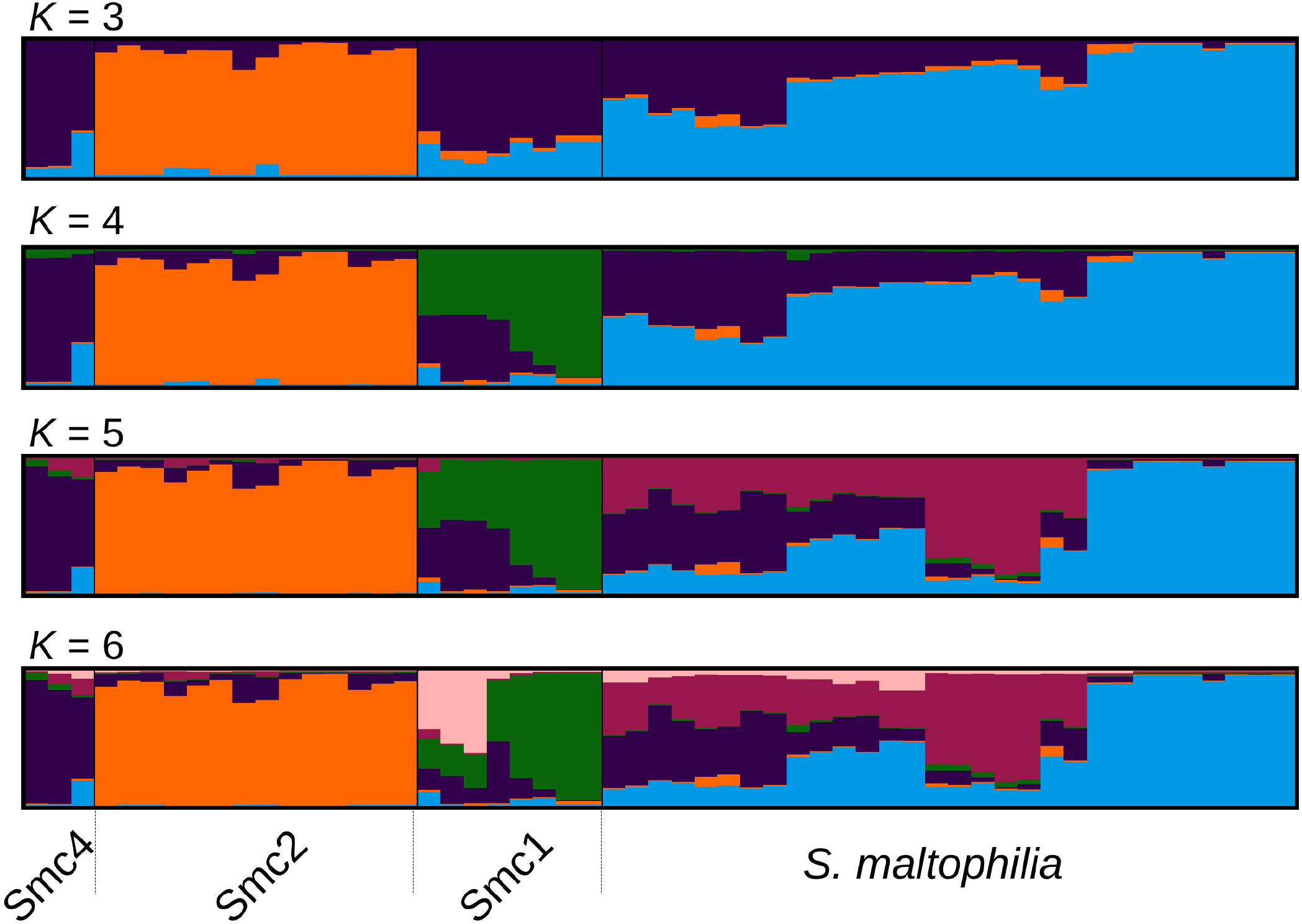
Optimally aligned STRUCTURE barplots for 20 replicate runs executed for *K* = 3 to *K* = 6 generated with CLUMPAK, showing the population genetic structure of the Mexican isolates from the Smc classified as *S. maltophilia* (Smalt) Smc1 (Sm1) and Smc2 (Sm2). The Sm4 cluster corresponds to Smc4 sequences from pubmlst.org, included in the analysis as an outgroup to the Mexican Smc strains (see Fig. 1B).

### 3.4 Population genetic structure analysis of Mexican environmental *S. maltophilia* complex isolates

In order to challenge the results of the multispecies coalescent (MSC)-based species delimitations presented in the previous section, we performed a Bayesian population clustering analysis on all Mexican isolates grouped within the Smc clade (Fig. 1B) using STRUCTURE (see methods). We included also the Smc4 lineage, identified among pubmlst.org sequences, as an outgroup control. Figure 2 shows the optimally aligned STRUCTURE barplots for the 20 replicate runs made for the indicated *K* values, depicting the ancestry proportions of each individual. Evanno’s delta-*K* method estimated an optimal number of 3 clusters, while Pritchard’s method suggested an optimal *K* = 6. This analysis uncovered a strong population subdivision of the Mexican Smc isolates, which is consistent with the MSC-based species delimitation (Fig. 1B). Given the independent evidence from the MSC analysis, we favor a conservative *K* = 4, as it consistently resolves the same 4 clusters classified as distinct species by the phylogenetic approach (*S. maltophilia*, Smc1, Smc2 and Smc4). Detailed inspection of the barplots reveals that already at *K* = 4 an important substructure exists within the *S. maltophilia* and Smc1 lineages. At *K* = 6 the Mexican *S. maltophilia* population gets subdivided into three clusters, while Smc1 is split into two subgroups. Clear evidences of admixture exist in both clades, suggesting that gene flow and recombination might be at play in these clusters.

### 3.5 McDonald-Kreitman (MK) neutrality test, genetic differentiation and gene flow estimates between pairs of environmental lineages of the *S. maltophilia* complex

To determine the statistical support of the major clusters revealed by STRUCTURE we computed the *K*^*^_ST_ index of population genetic differentiation (Hudson et al., 1992) between them. The results presented in Table 3 indicate that the Smc1, Smc2 and *S. maltophilia* lineages are strongly differentiated (*p* < 0.001) based on the *K*^*^_ST_ index, with multiple fixed differences between populations (range 9 – 45) and mean population divergences (*D*xy) > 4.5%. The high *F*_ST_ fixation indices (range 0.43 – 0.59) further denote very strong population differentiation. This is consistent with the low numbers of effective migrants per generation estimated (*Nm* range 0.34 – 0.66), indicating limited genetic flux between these lineages. We applied the MK test (McDonald and Kreitman, 1991) and computed the “neutrality index” (*NI*) (Rand and Kann, 1996) to test if the observed polymorphisms between pairs of lineages have evolved by the accumulation of neutral mutations by random drift, the fixation of adaptive mutations by selection, or a mixture of the two. As shown in Table 3, the *G* and *NI* indices for pairwise species comparisons indicate that fixed differences between species are due to nonsynonymous differences more often than expected, suggesting that positive selection may be driving divergence of Smc1 and *S. maltophilia* from Smc2. This signal is not significant (*p* = 0.077) for the Smc1 and *S. maltophilia* comparison.

**Table 3.** McDonald-Kreitman (MK) neutrality tests, genetic differentiation and gene flow estimates between environmental isolates of the Smc1, Smc2 and *S. maltophilia* (Smal) lineages of the *S. maltophilia* complex recovered from Mexican rivers based on the concatenated dataset (3591 sites). *D*xy is the interpopulation genetic distance. *K*^*^_ST_ is Hudson’s index of population genetic differentiation. *F*_ST_ is the fixation index. Nm represents the number of migrants per generation. *NI* is the neutrality index and *G* is the likelihood ratio or *G*-test of independence.

| Lineages | No. Fixed diffs. | $D_{xy}$ | $K_{ST}^*$ | $F_{ST}$ | $N_m$ | $NI$ | $G$ | $p$-value |
| --- | --- | --- | --- | --- | --- | --- | --- | --- |
| Smc1-Smc2 | 45 | 0.04616 | 0.13791*** | 0.57819 | 0.36 | 0.328 | 6.709 | 0.00959** |
| Smc1-Smal | 9 | 0.04542 | 0.06654*** | 0.43022 | 0.66 | 0.247 | 3.133 | 0.07672 |
| Smc2-Smal | 28 | 0.04924 | 0.09508*** | 0.59384 | 0.34 | 0.312 | 6.020 | 0.02515* |

<sup>a</sup> The significance of the estimated statistic was computed using the permutation test with 10000 iterations.
<sup>b</sup> $p$ -values for the $G$ -test.

### 3.6 Comparative analysis of DNA polymorphisms and recombination rates across the *S. maltophilia* complex (Smc) and *S. terrae* lineages

Table 4 presents basic descriptive statistics of DNA polymorphisms, neutrality and population growth tests computed for the Mexican populations/genospecies with > = 10 isolates (Smc1, Smc2, *S. maltophilia* and *S. terrae*. Based on their average nucleotide diversity per site (*π*) values, the lineages sort in the following decreasing order of diversity: *S. maltophilia* > Smc1 > *S. terrae* > Smc2. The high *π* values, high numbers of haplotypes (*h*) and the high haplotype diversity values (*Hd*) consistently reveal that species in the genus comprise notoriously diverse gene pools. Tajima’s *D* values are all negative, but non-significant, suggesting that the loci are either under purifying selection or that populations are undergoing expansions. We used the *R*_2_ population growth test statistic (Ramos-Onsins and Rozas, 2002) to test the null hypothesis of constant population size. Since all *p*-values were > 0.1, there is no evidence of population expansion.

**Table 4.** Descriptive statistics of DNA polymorphisms, neutrality and population growth tests for environmental isolates of the Smc1, Smc2, *S. maltophilia* (Smal) and *S. terrae* (Sterr) lineages recovered from Mexican rivers based on the concatenated dataset (3591 sites). The *p*-values for the *R_2_* statistic were estimated using 10000 coalescent simulations assuming an intermediate level of recombination.

| Species | No. Seqs | $S$ | $Eta$ | $\pi$ | $h/Hd$ | $Theta / site$ | $Tajima's D$ | $R_2$ | $p$-value |
| --- | --- | --- | --- | --- | --- | --- | --- | --- | --- |
| Smc1 | 11 | 258 | 280 | 0.02502 | 9/0.945 | 0.02662 | -0.29156 (NS) | 0.1390 | 0.45 |
| Smc2 | 15 | 173 | 185 | 0.01393 | 14/0.990 | 0.01584 | -0.45427 (NS) | 0.1199 | 0.19 |
| Smal | 51 | 500 | 560 | 0.02652 | 33/0.929 | 0.03470 | -0.85761 (NS) | 0.0903 | 0.17 |
| Sterr | 10 | 318 | 328 | 0.02388 | 7/0.067 | 0.04455 | -1.30647 (NS) | 0.1482 | 0.49 |

Table 5 shows the estimates for *R*/*theta*, the ratio between the mean number of recombination to mutation events. The ratios are > 1 (except for *S. terrae*) and for Smc2 and *S. maltophilia* the recombination events are estimated to be almost twice and nearly three times the number of mutation events, respectively. This indicates that homologous recombination events introduce significantly more polymorphisms into the *Stenotrophomonas* genomes than point mutations. The inverse mean DNA import length estimates (1/*delta*) suggest that on average rather long sequence stretches are affected by recombination (range 375–1472 nt), with a considerable mean sequence divergence (range 0.009–0.065).

**Table 5.** Recombination estimates for environmental isolates of the Smc1, Smc2, *S. maltophilia* (Smal) and *S. terrae* (Sterr) lineages recovered from Mexican rivers based on the concatenated dataset (3591 sites). The figures indicate the posterior mean and their variances are shown in parentheses.

| Lineages | $R/\theta$ | $1/\delta$ | $nu$ |
| --- | --- | --- | --- |
| Smc1 | 1.10774 (0.07076) | 0.0006789 (3.213e-08) | 0.0152529 (7.570e-07) |
| Smc2 | 1.89989 (0.12028) | 0.0009080 (3.121e-08) | 0.00864951 (3.081e-7) |
| Smal | 2.95326 (0.10044) | 0.0010379 (1.302e-08) | 0.0129774 (1.648e-7) |
| Sterr | 0.87233 (0.05113) | 0.0021663 (3.443e-07) | 0.0650391 (1.2279e-5) |

### 3.7 High diversity of novel STs among the Mexican Smc isolates

The allele numbers and STs were determined for each of the 77 isolates in the Smc (Smc1 = 11, Smc2 = 15 and *S. maltophilia* = 51) recovered from Mexican rivers, by comparing them with the corresponding 177 STs and associated alleles downloaded from pubmlst.org (as of Nov. 18^th^, 2016) using an in house Perl script. A high diversity of new alleles was discovered, as summarized in Table S4. Only the ST139, displayed by the Mexican *S. maltophilia* isolate ESTM1D_MKCAZ16_6, was previously reported in pubmlst.org, highlighting the novelty of the genotypes recovered in this study. A single entry is currently found in pubmlst.org for ST139, which corresponds to the Spanish isolate S3149-CT11 (id 220), recovered in 2007 from a surgical wound of a patient treated in an hospital from Barcelona. The Mexican Smc strains add 56 new STs (numbers 178 – 233) to those reported in the pubmlst.org database, with ST219 being the most prevalent one (see supplementary Tables S3, S4 and Figure S10), shared by 9 isolates recovered from the sediments of the highly contaminated TEX site (Table 1), both on MK and NAA media.

### 3.8 Multivariate association mapping of species, antimicrobial resistance phenotypes and habitat preferences by multiple correspondence analysis (MCA)

We used MCA to visualize the associations between the antibiotic resistance profiles, β-lactamase production phenotypes (Fig. S11), habitat preferences and species assignations (Figs. 1A and 1B) made for the Mexican *S. maltophilia*, Smc1, Smc2 and *S. terrae* isolates (all with > 10 isolates/species). Figure 3 depicts the MCA factor-individual biplot resulting from the analysis of 17 active variables and 4 supplementary categorical variables, listed in the figure caption. The clouds of individuals for each species were hidden (visible in Fig. S12) to avoid over-plotting, but the 95% confidence intervals (CIs) for species are shown as color-coded ellipses. The first two dimensions explain 45.3% of the variance, the first dimension accounting for > 3.8 times the variability explained by the second and following ones, as shown in the screeplot presented in Fig. S13A. The variable plot depicted in Fig. S13B reveals that the active variable species is most correlated one with the two first two dimensions, indicating that their resistance profiles and β-lactamase expression phenotypes are distinct. The variables GM, KA, MER, FEP, TET, ATM (abbreviations defined in the legend to Fig. 3), β-lactamase expression and the MDR condition are strongly correlated with the first component, while IMP and SM are the variables most strongly associated with the second dimension (Fig. S13B). Figure 3 shows that the *S. maltophilia* (malt) strains form a distinct and independent cloud that is characterized by a very strong association with a resistance status to the following antibiotics: CAZ, CAZ.CLA, GM, FEP, GM, KA and SM. It is also strongly associated with the MDR condition and metallo-β-lactamase production. The latter are the most-strongly contributing variables for the delimitation of this group, as depicted in the variable-categories MCA map presented in Fig. S14. *S. maltophilia* shows a preference for the sediments of contaminated sites. The resistance phenotypes and habitat preferences of the Smc1 and Smc2 lineages largely overlap, those of *S. terrae* being more differentiated, but partially overlapping with the 95% CI ellipse for Smc2. The Smc1 and Smc2 lineages are strongly associated with non MDR, aminoglycoside, P.T. and Tm.Cb sensitivity, showing a preference for the water column of clean or moderately contaminated sites (Fig 3 and Fig S13). The *S. terrae* isolates are distinctly and strongly associated with carbapenem and ATM sensitivity (Fig 3 and Fig S14). The statistical significance of these antibiotic resistance and habitat preference patterns will be formally tested in the next two sections, respectively.

**Figure 3.**
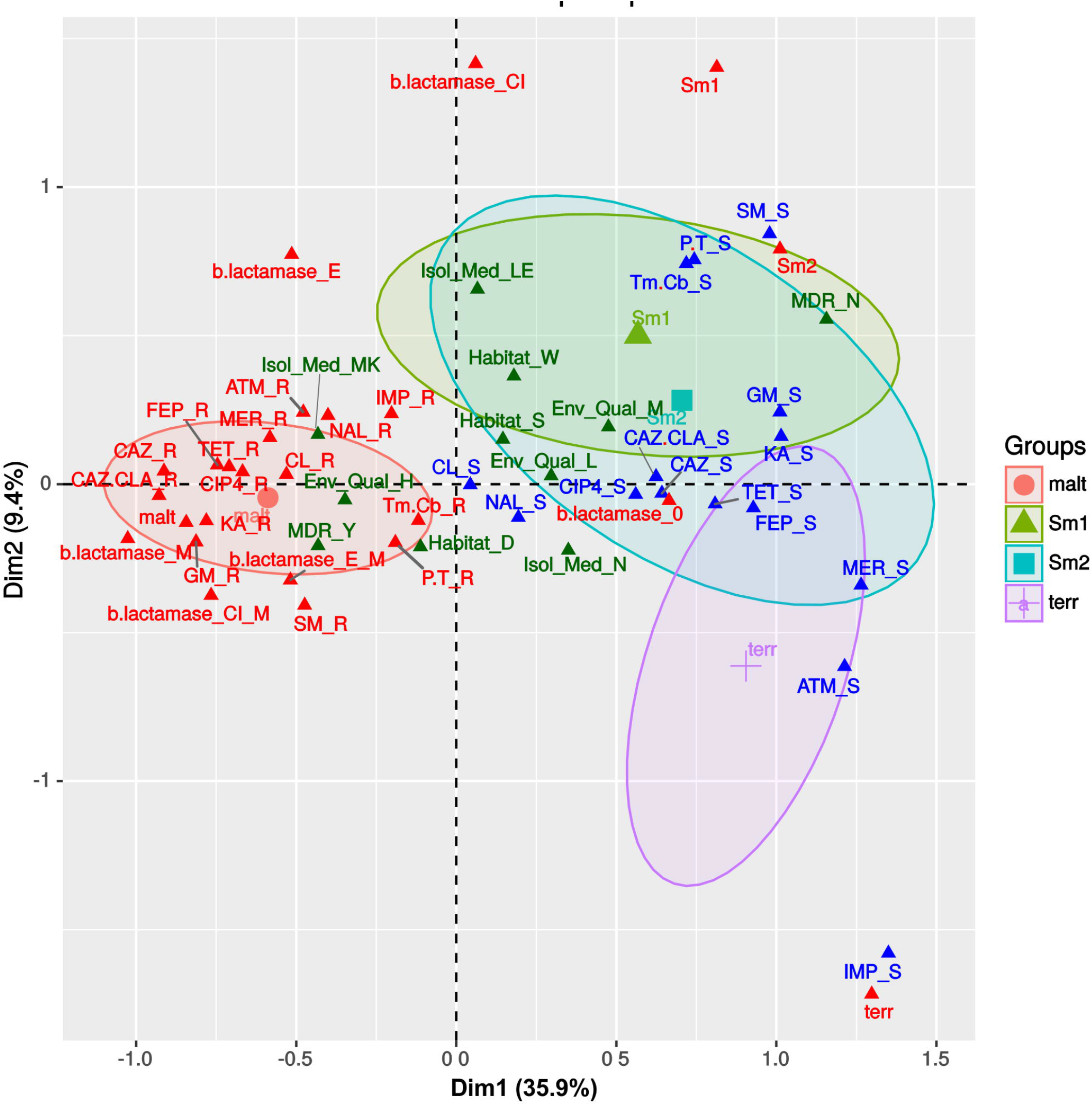
Multivariate correspondence analysis (MCA) factor –individuals biplot map, summarizing the associations between antibiotic resistance profiles, β-lactamase production phenotypes and species assignations. The state of 17 active variables (IMP=imipenem, MER=meropenem, CAZ=ceftazidime, CAZ.CLA=ceftazidime/clavulanate, FEP=cefepime, ATM=aztreonam, b.lactamase, species, Tm.Cb=trimethoprim+carbenicillin, CL=chloramphenicol, SM=streptomycin, GM=gentamicin, KA=kanamycin, P.T=piperacillin/tazobactam, NAL=nalidixic acid, CIP4=ciprofloxacin, TET=tetracycline) and 4 supplementary categorical variables [Habitat, Env_Qual (environmental quality), Isol_Med (isolation medium), MDR] are shown, sorted along the first two dimensions that together explain 45.3% of the total variance. Some variables like CTX4 (cefotaxime) were excluded, due to lack of variability in the observed states. Species name abbreviations are as follows: malt = *S. maltophilia*; Sm1 = Smc1; Sm2 = Smc2; terr = *S. terrae*.

### 3.9 Only *S. maltophilia* is truly multidrug-resistant (MDR) and isolates from polluted sites express resistance to more antibiotic families

We performed one-way ANOVA analyses to evaluate differences *i*) in the mean number of resistances to individual antibiotics (NumR), *ii*) in the mean number of distinct drug families (NumFam) across species, and *iii*) to determine whether *S. maltophilia* isolates recovered from high and low pollution sites have the same NumR and NumFam. Figures 4A, 4B, 4E and 4F display violin and boxplots for the raw count data, which revealed skewed, non-normal distributions, with a few outliers. Key assumptions (homoscedasticity and normality) made by standard one-way ANOVA were formally tested (Tables S5, and S6), which confirmed multiple cases of highly significant departures from normality. Consequently, we performed robust one-way ANOVA (Wilcox, 2016) using trimmed means (*tr* = .2) and bootstrap (*nboot* = 2000) to simulate the distribution of the trimmed sample means and compute the appropriate critical values for the confidence intervals (95% CIs). Figs. 4C and 4D depict mean plots with 95% CIs for the NumR and NumFam, respectively, across the four species with > 10 isolates. These clearly show that *S. maltophilia* strains express significantly higher mean_(*tr* =.2)_ NumR (12.63) and NumFam (4.66) than the other three species, only *S. maltophilia* being truly MDR. Highly significant results for both the NumR [*F_t_* = 176.3447, *p* = 0. Variance explained (*σ^2^*) = 0.821; effect size (*e.s*.) = .906] and NumFam (*F_t_* =32.2755, *p* = 0; *σ^2^* = .511; *e.s*. = .715) were obtained, thus rejecting the null hypothesis of equal trimmed means for both variables across species, and revealing huge effect sizes (>.8). A Wilcox robust *post-hoc* test of the trimmed mean NumR and NumFam comparisons across pairs of species confirms that all those involving *S. maltophilia* are very highly significant (*p* = 0; Figs. S15A and S15B, Tables S7 and S8), as indicated by asterisks on the mean_(*tr* =.2)_ plots shown in Figs. 4C and 4D. Sample sizes for *S. maltophilia* isolates recovered from sites with high and low pollution were large enough (Fig. 4E) to test the hypothesis of equal NumR and NumFam conditional on pollution level. We used the robust *yuenbt*(*tr* = .2, *nboot* = 998) method for independent mean comparisons (Wilcox, 2016). The tests for NumR [*T_y_* = 2.1964, *p* = 0.038; mean_(*tr* =.2)_ difference = 1.233, *CI*_95%_ = (0.0663,2.4004), *e.s*. = .43] and for NumFam [*T_y_* = 2.6951, *p* = 0.022; mean_(*tr* =.2)_ difference = 0.95, *CI*_95%_ = (0.1636, 1.7364), *e.s*. = .5], indicating that both variables have significantly higher values in the high-pollution sites (Figs. 4G and 4H). However, given that the 95% *CIs* of the mean NumR comparsion overlap (Fig. 4G), and that a high number of tied (repeated) values occur (Fig. 4E), we run also Wilcox’s robust percentile bootstrap method for comparing medians [*medpb2*(*nboot* = 2000)], which provides good control over the probability of Type I error when tied values occur. For NumR the test could not reject the null of equal medians [*M\** =1, *p* = .094, *CI*_95%_ = (0, 2.5)], but was significant for NumFam [*M\** =1, *p* = .0495, *CI*_95%_ = (0, 2)].

**Figure 4.**
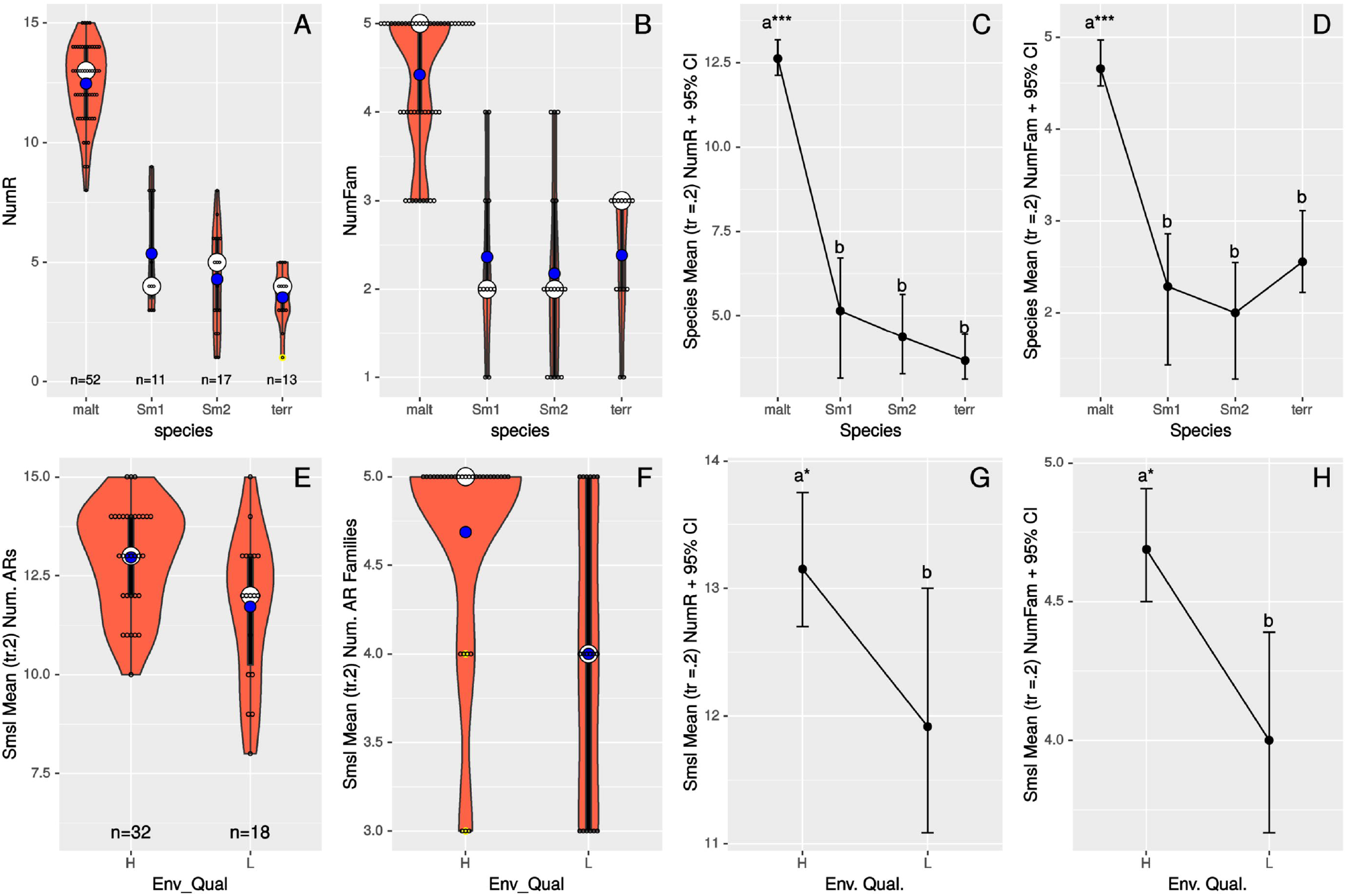
Panels A, B, E and F display violin and boxplots for the raw count data of number of resistances to individual antimicrobials (NumR) and distinct drug families (NumFam) for Mexican environmental isolates. The big white dot shows the median and the smaller blue one the mean of the distribution of individual observations, represented as small open circles. Yellow dots indicate outlier data points. Panels C, D, G and H show meanplots for NumR and NumFam (see labels on the ordinates) for 20% trimmed means and 95% confidence intervals estimated by non-parametric bootstrap (*nboot* = 1000). Upper row panels (A-D) correspond to analyses for the Mexican isolates classified in the four species/lineages indicated on the x-axes. Lower panels (E-H) correspond to Mexican *S. maltophilia* isolates recovered from high and low pollution sites, based on the criteria indicated on Table 1. The 4 *S. maltophilia* isolates recovered from sites with intermediate contamination level were excluded, as only populations of organisms with > 10 isolates were considered. Species codes are as defined in Fig. 3.

### 3.10 *Stenotrophomonas* species differ in their β-lactamase expression patterns and only *S. maltophilia* strains express metallo-β-lactamases

Association plots (Fig. 5) revealed a very highly significant association (*p* ≈ 0) between species and type of β-lactamases expressed (Fig. S11). Metallo-β-lactamase expression was significantly and exclusively associated with *S. maltophilia* isolates. Most isolates from the Smc1 and Smc2 lineages did not express any kind of β-lactamase, although expression of extended spectrum of β-lactamases could be detected in a few isolates of these species. No β-lactamase expression was detected in *S. terrae* isolates.

**Figure 5.**
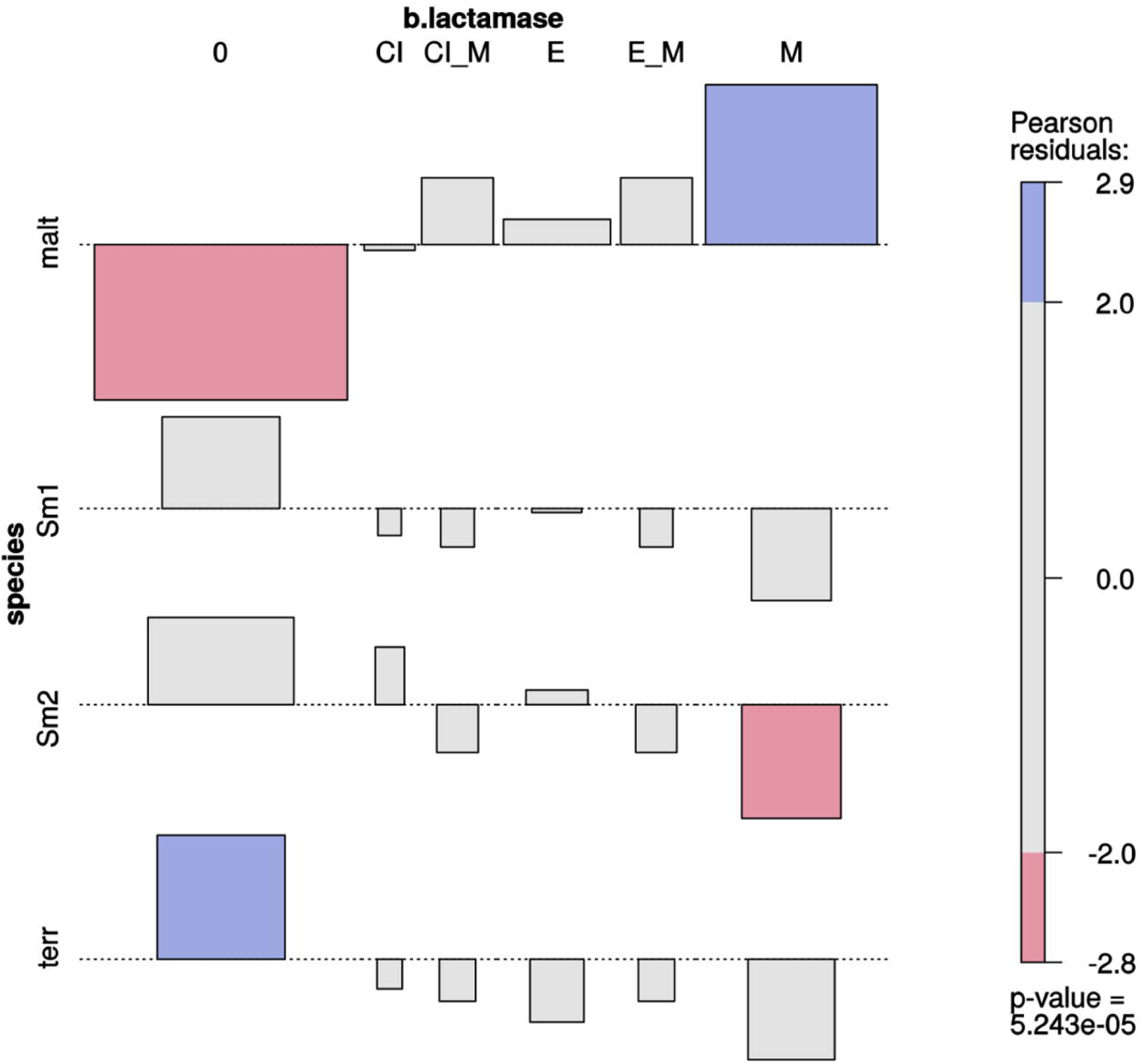
Two-way association plot for the categorical variables species and β-lactamase production phenotype and *Stenotrophomonas* species containing > 10 Mexican environmental isolates. The bars on the plot represent the Pearson residuals, the color code and the height of the bars denote the significance level and magnitude of the residuals, and their widths are proportional to the sample size. The β-lactamase codes are as follows: 0 = no β-lactamase activity detected; CI = clavulanate-inducible class A or class C (AmpC) cephalosporinase; CI_M = clavulanate-inducible cephalosporinase plus metallo β-lactamase (MBL); E = extended-spectrum β-lactamase (ESBL); E_M = ESBL plus MBL; M = MBL. Species codes are as defined in Fig. 3.

### 3.11 The prevalence of environmental *Stenotrophomonas* species recovered from Mexican rivers is significantly associated with habitat and pollution level

We performed a multi-way association analysis to test the null hypothesis that *Stenotrophomonas* species prevalence is independent of isolation habitat (sediment *vs*. water column), the pollution level of the sampling site (Table 1) and isolation medium (Fig. 6). The test strongly rejects the null hypothesis (*p* < .00001). *S. maltophilia* was mainly recovered on MK plates, being significantly associated with polluted sediments. The Smc1 lineage displayed a moderately significant association with clean water columns, although some isolates could also be recovered from contaminated sediments using NAA, which is consistent with the MCA results presented in Fig. 3. In contrast, Smc2 isolates were very significantly overrepresented in the water columns of clean rivers and underrepresented in sediments, suggesting a high level of ecological specialization. *S. terrae* isolates were mainly recovered on oligotrophic NAA plates from the sediments of clean sites (Table 1).

**Figure 6.**
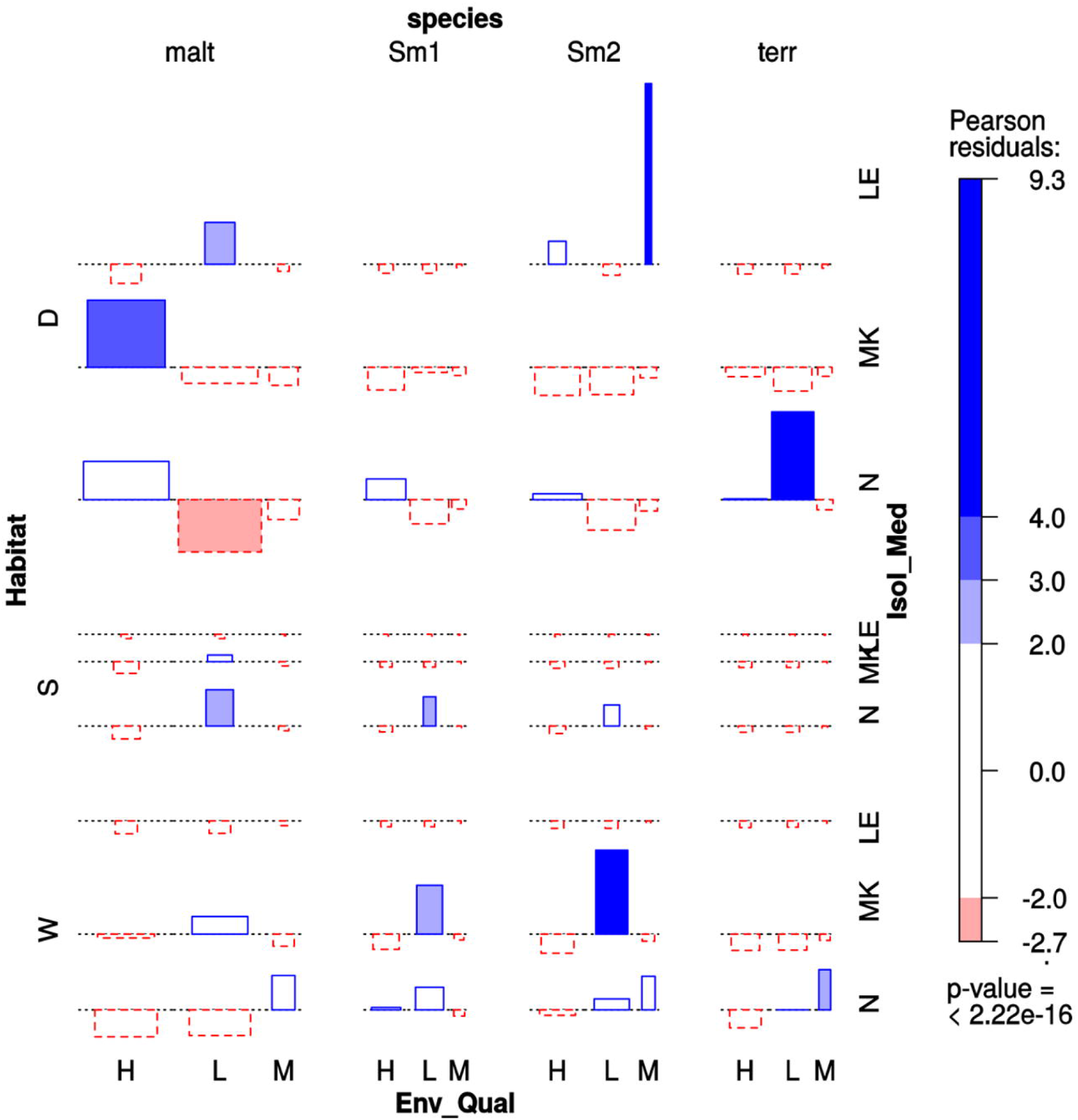
Four-way association plot showing the results of multiway-chi-square analysis for the categorical variables species (abbreviations as defined in Fig. 3), habitat (W = water column; S = flooded soil; D = sediment), isolation medium (N = NAA; MK = MacConkey; LE = Leed’s Medium) and pollution level (H = high; L = low; M = intermediate, based on counts of thermotolerant coliforms and *E. coli*, as defined in Table 1), using Friendly’s residual coloring scheme to highlight the significant associations. Species codes are as defined in Fig. 3. The bars on the plot represent the Pearson residuals, the color code and the height of the bars denote the significance level and magnitude of the residuals, and their widths are proportional to the sample size.

## 4. Discussion

In this study we demonstrate the power of complementary phylogenetic and population genetics approaches to delimit genetically and ecologically coherent species among a diverse collection of environmental isolates of the genus *Stenotrophomonas*. Importantly, this is done based exclusively on molecular evolutionary criteria (Vinuesa et al., 2005b), without using any arbitrary sequence or phenetic similarity cut-off values, as embraced by the standard polyphasic approach that dominates current bacterial taxonomic practice (Kämpfer and Glaeser, 2012). The robustness of the species delimitations proposed here are supported by the statistically significant associations they exhibit with distinct habitat preferences, antibiotic resistance profiles, MDR status and β-lactamase expression phenotypes.

To our knowledge, this is the first study that used the multispecies coalescent (MSC) model (Rannala and Yang, 2003;Edwards et al., 2007;Degnan and Rosenberg, 2009) coupled with Bayes factor (BF) analyses (Kass and Raftery, 1995) for microbial species delimitation. The MSC model has the virtue of relaxing the implicit assumption made by the concatenation approach that the phylogeny of each gene is shared and equal to that of the species tree. Although this assumption is problematic, the concatenation approach is the current standard in microbial multilocus sequence analysis (Gevers et al., 2005;Vinuesa, 2010;Glaeser and Kampfer, 2015), including phylogenomics (Rokas et al., 2003;Wu and Eisen, 2008). It has been shown that phylogenetic estimates from concatenated datasets under coalescence are inconsistent (Kubatko and Degnan, 2007;Song et al., 2012), that is, converge to wrong solutions with higher probability as the number of concatenated loci increases. However, the impact of this inconsistency still needs to be thoroughly evaluated with clonally multiplying microbial organisms experiencing different rates of recombination (Hedge and Wilson, 2014). In our analyses, the topology of the ML phylogeny inferred from the concatenated dataset (Fig.1) is largely congruent with the Bayesian species tree inferred under the MSC (Fig S8). It is worth noting that the numbers on the branches in this type of species tree denote the estimated population sizes. That of *S. maltophilia* is about 1 order of magnitude larger than the population size estimates for the Smc1 and Smc2 genospecies (Fig. S8), which reflects the genetic heterogeneity of strains grouped in the *S. maltophilia* lineage, which includes the majority of the Mexican isolates, along with reference strains from the pubmlst.org and genome databases, isolated across the globe. Lumping this heterogeneous set of recombining sub-lineages into a single species results in coalescent events higher up in the species tree than those observed for the Smc1 and Smc2 lineages. This is consistent with the marked internal structure revealed by the Bayesian structure analysis within *S. maltophilia* (Fig. 2), which suggests that additional cryptic species may be found within the *S. maltophilia* lineage. Patil and colleagues recently proposed that the type strains of *S. africana*, *P. beteli* and *P. hibiscicola*, which are phylogenetically placed within the *S. maltophilia* clade (Fig. 1A), and have been reclassified as *S. maltophilia*, actually represent distinct species, based on their estimates of genomic average nucleotide identity values < 94% (Patil et al., 2016). In our view, these clusters represent incipient species that are still capable of recombining between them, as suggested by the admixture found in the STRUCTURE barplots (Fig. 2) and by our estimates for recombination within *S. maltophilia* (Table 5). Further investigations involving comparative and population genomics are required to identify clear signatures of speciation within the *S. maltophilia* sub-lineages, including “speciation genes and islands” (Shapiro et al., 2016). Despite the conservative approach taken in this study, the BF analysis provides statistical support in favor of splitting the *S. maltophilia* complex as currently defined in pubmlst.org into the following 5 broad evolutionary lineages: *S. maltophilia* and the genospecies Smc1 to Smc4. Overwhelming support (ln-BF > 5) was also obtained indicating that genogroup #10 (Vasileuskaya-Schulz et al., 2011), the sister clade of *S. rhizophila* (genogroup #8), constitutes an independent, non-described species (Table 2). This is of practical importance from a biotechnological perspective because it has been argued that plant-associated *S. rhizophila* strains (Vasileuskaya-Schulz’s genogroup #8) can be safely and easily separated from *S. maltophilia* pathogens in clade IIb (Berg and Martinez, 2015) based on 16S rRNA gene sequences and *ggpS* and *smeD* PCR-based typing (Ribbeck-Busch et al., 2005). However, it would be important to define differences of the former with strains in the sister genogroup #10, which holds both rape rhizosphere and human blood and tigh bone infection isolates. In summary, our MSC-BF analysis provided strong evidence for the existence of 5 new species in the analyzed dataset.

Considering that the MSC model implemented in *BEAST 2 was not specifically developed for bacteria, and given that this model has been put under criticism due to detectable model misspecification when tested on diverse empirical animal and plant datasets using posterior predictive simulations (Reid et al., 2014), it was important to evaluate the robustness of the Bayesian species delimitations with independent methods. We challenged the proposed species borders within the Smc by performing well-established population genetic analyses on our collection of environmental isolates from the sister lineages *S. maltophilia*, Smc1 and Smc2. We focused on detecting population genetic structure, estimating gene flow between the phylogenetically defined species, and identifying signatures of selection. Such data and evidence are predicted by current ecological models of bacterial speciation to reflect speciation events (Vinuesa et al., 2005b; Koeppel et al., 2008;Vos, 2011;Cadillo-Quiroz et al., 2012;Shapiro and Polz, 2014).

Current models of bacterial speciation suggest that groups of closely related strains that display some degree of resource partitioning, and consequently occupy different niches, will be affected by independent selective sweeps caused by the gain of a beneficial gene either by horizontal transfer or by adaptive mutation. These may sweep to fixation if recombination levels are low in relation to selection coefficients, or form so called “speciation islands or continents” in the genomes of highly recombining populations (Cadillo-Quiroz et al., 2012;Shapiro et al., 2012). Such (sympatric) populations are predicted to be discernable as sequence clusters that diverge from each other because of the fixation of different adaptive mutations. This leads to the formation of independent genetically and ecologically coherent units as gene flow between them gradually drops as they diverge by means of natural selection (Vos, 2011;Shapiro and Polz, 2014). We could show that the Mexican *S. maltophilia*, Smc1 and Smc2 lineages satisfy these predictions. The *K^*^*_ST_ test statistic (Hudson et al., 1992) detected highly significant genetic differentiation between all pairs of these sympatric lineages based on DNA polymorphisms (Table 3). This is consistent with the results from the STRUCTURE analysis. Conversely, the number of migrants between these lineages was negligible, evidencing low levels of gene flow between them. Additionally, the “neutrality index” (*NI*), and the results of the MK tests suggest that positive selection, rather than drift, is the force promoting divergence between the lineages, which is in line with predictions from the adaptive divergence model (Vos, 2011). However, the latter interpretation needs to be considered cautiously, as the high relative rate of between-species non-synonymous substitutions observed could also be generated by within-species purifying selection to eliminate slightly deleterious mutations (Hughes, 2005;Hughes et al., 2008). The latter interpretation is consistent with the observed negative, but not significant Tajima’s *D* values (Table 4). These are not likely to reflect a population expansion, given the non-significant *p*-values of the powerful *R*_2_ statistic for population growth, which is well suited for small sample sizes such as those of the Smc1 and Smc2 lineages (Ramos-Onsins and Rozas, 2002). We could show that recombination is an important force, providing genetic cohesion to these lineages, with *Rho/theta* estimates ranging from 1.11 in Smc1 to nearly 3 in *S. maltophilia*. Since recombination events are only detectable when a tract of multiple polymorphisms are introduced in a population, it is clear that most of the observed polymorphisms within the analyzed populations originate from recombination rather than point mutations. The high recombination levels detected within the *S. maltophilia* lineage suggests that speciation within this group is an ongoing, possibly not yet finished process, along a “spectrum” of speciation, resulting in “fuzzy” borders between the sub-lineages (Hanage et al., 2005;Shapiro et al., 2016). However, these can be already detected as phylogenetic and STRUCTURE clusters, even with the limited resolving power provided by the 7 gene MLST scheme used.

As predicted by the ecological speciation models recently developed for bacteria (Koeppel et al., 2008;Vos, 2011;Shapiro and Polz, 2014), the marked genetic differentiation detected between the sympatric *S. maltophilia*, Smc1 and Smc2 environmental populations is significantly associated with different habitat preferences, antibiotic susceptibility profiles and β-lactamase expression phenotypes. These attributes strongly suggest that these lineages have differentiated ecological niches. The significant differences in habitat preferences could provide some micro-geographic separation between the populations coexisting in the same river, which might partly explain the reduced gene flow measured between them, despite of being sister lineages, contributing to their genetic differentiation. Similar patterns have been reported for other aquatic microbes such as *Desulfolobus* (Oakley et al., 2010), *Exiguobacterium* (Rebollar et al., 2012) and *Vibrio* (Shapiro et al., 2012;Friedman et al., 2013). Consequently, our results support the growing body of evidence pointing to niche partitioning as a major factor promoting evolutionary divergence between closely related sympatric prokaryotic populations, even when they exhibit high levels of recombination (Shapiro and Polz, 2014).

As noted before, *S. maltophilia* is well-known as an emergent opportunistic multidrug-resistant (MDR) nosocomial pathogen, causing increasing morbidity and mortality (Looney et al., 2009;Brooke, 2012). Comparative genomics and functional analyses have clearly established that the MDR or extensively drug-resistant (XDR) phenotype displayed by clinical isolates of this species is largely intrinsic, resulting from the expression of a combination of several types of efflux pumps (RND, MATE, MFS and ABC types) and diverse chromosomally-encoded antibiotic resistance genes [*aph (3’)-IIc*, *aac(6’)-lz* and Sm*qnr*], including the metallo-beta-lactamase *blaL1* and the inducible Ambler class A beta-lactamase *blaL2* (Crossman et al., 2008;Brooke, 2014;García-León et al., 2014;Sanchez, 2015;Youenou et al., 2015). However, contradictory results have been reported regarding the MDR status of environmental isolates of the Smc. For example, a recent ecological study of a large collection isolates classified as *S. maltophilia* recovered from diverse agricultural soils in France and Tunisa concluded that they display a high diversity of antibiotic resistance profiles, expressing resistance against 1 to 12 antibiotics, with clinical and manure isolates expressing the highest numbers (Deredjian et al., 2016). These isolates were vaguely classified as *S. maltophilia* based on growth on the selective VIA isolation medium (Kerr et al., 1996) and PCR detection of the *smeD* gene (Pinot et al., 2011). We argue that the large phenotypic variance observed in that and similar studies result from the lack of proper species delimitation. This cannot be achieved with such coarse typing methods, most likely resulting in the lumping of multiple species into *S. maltophilia*. In contrast, in the present study we found a very strong statistical association between the MDR condition and metallo-β-lactamase (MBL) production with the *S. maltophilia* lineage, whereas the sibling Smc1 and Smc2 genospecies were found to express on average resistance to < 3 antibiotic families (Figs. 4B and 4D and S13B), most strains not expressing any kind of β-lactamase, and none expressing MBLs (Fig. 5). Consequently, intrinsic MDR can only be assumed for the *S. maltophilia* strains of clinical and environmental origin.

In conclusion, the results presented here provide the first in depth and integrative molecular systematic, evolutionary genetic and ecological analysis of the genus *Stenotrophomonas*. The study demonstrates that both phylogenetic and population genetic approaches are necessary for robust delimitation of natural species borders in bacteria. Failure to properly delimit such lineages hinders downstream ecological and functional analysis of species. Comparative and population genomic studies are required to resolve pending issues regarding the speciation status of the sub-lineages within the *S. maltophilia*.

## 6. Author Contributions

All authors read and approved the manuscript. LEOS and PV conceived and designed the project. LEOS generated the collection of isolates, performed wet-lab experiments and analyzed resistance phenotypes. PV performed bioinformatics, statistical and evolutionary genetic analyses. PV wrote the manuscript.

## 8. Acknowledgements

We gratefully thank Dr. Eria Rebollar and Dr. Claudia Silva for their critical reading of the manuscript. Javier Rivera Campos is acknowledged for technical support with wet-lab experiments. Antonio Trujillo from CCG-UNAM is thanked for support with field work. Don Juan Alvear Gutiérrez and Ing.Norberto Bahena are gratefully acknowledged for supporting our sampling at the natural parks Los Sauces and Las Estacas, respectively. José Alfredo Hernández and the UATI at CCG-UNAM are acknowledged for support with Linux server administration. Dr. Jesús Silva Sánchez from the INSP in Cuernavaca, Mexico, is gratefully thanked for his support throughout the work, particularly regarding the interpretation of disk-diffusion assays and for providing laboratory reagents.

